# Consequences of Cancer on Zebrafish *Danio rerio*: Insights into Sex Determination, Sex Ratio, and Offspring Survival

**DOI:** 10.1101/2024.02.08.579516

**Authors:** Justine Boutry, Mathieu Douhard, Klara Asselin, Antoine M. Dujon, Jordan Meliani, Olivier De Backer, Delphine Nicolas, Aaron G. Schultz, Peter A. Biro, Christa Beckmann, Laura Fontenille, Karima Kissa, Beata Ujvari, Frédéric Thomas

## Abstract

Offspring sex ratio has been proposed as an indicator of the risk of developing certain cancers in humans, but offspring sex ratio may also be a consequence of the disease. In this study, we delve into this subject using the fish *Danio rerio* as a model system. First, we explore whether inducing skin cancer at an early stage of the host’s life (embryonic stage) has the potential to influence sex determination and/or sex-specific mortality. Second, we investigate whether the sex ratio in offspring produced by tumor-bearing adult females differs from that of healthy females. Third, we compare the survival (until sexual maturity) of offspring produced by cancerous and non-cancerous females. We found that skin cancer did not influence sex ratio in both experiments. However, consistent with previous studies on other model systems, the survival of offspring from cancerous females was higher, suggesting that diseased females allocate more resources to current reproductive efforts compared to their healthy counterparts. This study makes a significant contribution to our understanding of the ecological and evolutionary consequences of host-tumor interactions in animals.

## INTRODUCTION

While it is now widely acknowledged that numerous human activities promote cancerous pathologies in wildlife (Giraudeau et al. 2018)(Sepp et al. 2019)(Baines et al. 2021a), the exact magnitude of the resulting ecological and evolutionary consequences remains incompletely understood at present (Vittecoq et al. 2013)(Thomas et al. 2017). Simply predicting the accelerated disappearance of animals that have developed tumors in ecosystems is an oversimplified viewpoint that overlooks other potential repercussions. For instance, at the ecosystem level, the ecological impacts of differing susceptibility to cancer among species will vary based on the functional traits of the most affected species (e.g. predators, prey, keystone species (Hamede et al. 2020)(Cédric Perret, Cindy Gidoin, Beata Ujvari)). Phenotypic changes induced by oncogenic processes in hosts also have the potential to alter various interactions between affected organisms and other species within the ecosystem. For example, tumor-bearing hydra exhibit modified interactions with other species compared to their healthy counterparts – they capture more prey, are more susceptible to predation, and experience heavier colonization by commensal ciliates (Boutry et al. 2022a) (see also (Duneau and Buchon 2022)). At the individual and species levels, Dujon and colleagues (submitted) have suggested that effects at different time scales should be considered in the case of sudden and chronic exposure to mutagenic substances. Initially, it is expected that organisms will over-activate their anticancer defenses, significantly impacting them because these defenses are energetically costly and may force energetic trade-offs (Dujon et al. 2022)(Thomas et al. 2019)(Hiske Klaassen, Sophie Tissot, Jordan Meliani, Justine Boutry, Anna Miltiadous, Peter A. Biro, David J. Mitchell, Beata Ujvari, Aaron Schultz, Frédéric Thomas 2024)(Peter A. Biro, Frédéric Thomas, Beata Ujvari). It is also anticipated that affected species may progressively alter their life history traits, for example favoring earlier investment in reproduction (e.g. (Arnal et al. 2017)(Boutry et al. 2022b)(Jones et al. 2008)). In the long term, it is possible that natural selection will favor the development of more powerful anticancer defenses, such as duplications of tumor suppressor genes (e.g. (Sulak et al. 2016)(Trivedi et al. 2023)). Thus, the consequences of anthropogenically induced oncogenic activities on wildlife are diverse and complex, warranting further in-depth research, especially in aquatic ecosystems that are particularly prone to pollution (Häder et al. 2020).

The zebrafish (*Danio rerio*) is a widely used model for studying cancer due to its ability to develop cancer spontaneously after exposure to a mutagen or by transgenesis (Bambino and Chu 2017). Many tumors that *Danio* develop are highly homologous with certain human cancers at histological, proteomic and genetic levels (Kobar et al. 2021). In addition, research has shown that a mixture of organic pollutants common in today’s environments can affect development and behavior in zebrafish, highlighting the potential threats of these pollutants to zebrafish and, by extension, to human and animal populations (Bambino and Chu 2017). However, the extent to which the higher incidence of tumors could have population-wide repercussions through changes in life-history traits (Ujvari et al. 2016)(Boddy et al. 2015) such as sex ratios or reproductive investment has so far received little consideration.

In humans, offspring sex ratio has been proposed as an indicator of risk of developing certain cancers in both women and men. There is increasing evidence that reproductive hormones play a role in the process of sex ratio adjustment in mammals (Grant and Chamley 2010)(Merkling et al. 2018). Several review papers suggested that hormone profile at the time when children were conceived is responsible for both the offspring sex ratio and the incidence of cancer (James 2008)(James 2013). For instance, the high proportion of sons among women with pre-menopausal breast cancer may reflect the high levels of estrogen at conception (James 2006). Men who develop testicular cancer have, prior to the cancer diagnosis, a lower proportion of boys compared with the general population (James 2006), possibly due to low testosterone concentrations at the time of conception. However, data regarding whether the children were conceived before or after disease diagnosis is not sufficient to determine how these variables directly influence each other because oncogenic processes frequently exist at a sub-clinical level earlier in life (Thomas et al. 2018). In patients with multiple sclerosis, the offspring sex ratio biases occur after disease onset but the consequence of the disease and its treatment are particularly difficult to distinguish (James 1996). Thus, studies examining the consequence of cancer on sex allocation in model organisms are warranted.

The aim of this study was to provide one of the first experimental studies of the effects of cancer on biological traits relevant to evolutionary ecology. First, we examine the possibility that skin cancer occurring at a very early age of the host (i.e. embryonic) could affect sex-specific mortality and/or sex determination, which is partially influenced by the environment in zebrafish (Experiment 1). This bias may occur due to a higher mortality among male embryos developing cancer compared to female embryos (e.g. (Li et al. 2017), see also (Poulin 1996)(Wells 2000)(Lary and Paulozzi 2001) for general references). Because sex differences in juvenile mortality may influence the strength of selection for sex allocation (Cox and Calsbeek 2010), cancer may increase the likelihood of producing females rather than males. Next, we investigated whether adult females bearing tumors, compared to healthy females, have a sex ratio bias in their brood (Experiment 2). This hypothesis was motivated in part by the influence of toxic substances or high population density on the sex ratio of subsequent generations through epigenetic mechanisms in zebrafish (Noëmie et al. 2022)(Pierron et al. 2021). A biased sex ratio favoring males could also occur due to the expectation that females with health issues will produce lower-quality eggs. Regarding the overall survival of offspring in experiment 2, a first potential prediction is that, being the progeny of mothers in poor health, these offspring start life with a disadvantage, potentially reducing their chances of survival. On the other hand, it cannot be dismissed that females with cancer, as observed in other species, allocate more resources to their immediate reproductive events (see references above). This, as a terminal investment in reproduction (Clutton-Brock 1984), may actually contribute to the enhanced survival of their offspring.

## MATERIAL AND METHODS

### Zebrafish sex determination and differentiation

All zebrafish larvae develop ovary-like tissue between 13-25 days postfertilization (Kossack and Draper 2019). Between 20- and 25-days post-fertilization, the early-stage oocytes for individuals that were determined as female continue to mature, while for individuals that were determined as males their early-stage oocytes undergo apoptosis as the gonadal transition from a proto-ovary into a functional testis. Sexual maturation is not complete until zebrafish are 2.5-3 months old (Kimmel et al. 1995). The times listed above are approximative and can vary depending on rearing conditions such as temperature. The mechanism of sex determination in zebrafish is complex, but once determined, remains the same throughout life. While wild zebrafish have a genetic WZ system (see (Wilson et al. 2014)), this sex determinant appears to have been modified during domestication (Kossack and Draper 2019). In domesticated lines a polygenic system regulates sex, but the particular genes involved vary between the independently selected lines. It is also known that sex ratios in domesticated lines vary depending on certain environmental factors, such as temperature, pH, oxygen concentration and rearing density (Santos et al. 2017).

### Rearing protocol

Zebrafish were hatched, raised and monitored by Azelead fish facility (Montpellier, France) ©, experiments were conducted in three main experimental random batch (exp1; 12/11/2020 - 03/15/2021, Exp2; 04/19/2012 – 07/20/2021, Exp3; 09/28/2021 – 01/03/2022). Fish larvae from their 4th to 10th day were raised in Pétrie dishes of 10cm of diameter with a maximal density of 1 embryo/cm^2^ and fed dry food (75 µg, *Plantkovie ©, France*) four times daily. Juvenile fish between their 10th and 75th day were raised in 1L tanks at a maximal density of 50 fish and fed with a mixture of artemia nauplii (*Artemia salina, JBL©, Germany*) twice daily, and dry food (200 µg) four times a day. Adult fish older than 75 days were raised in 2,6L tanks and fed with both artemia nauplii *(Artemia salina, JBL©)* and dry food (300 µg) twice daily. Artemia nauplii were hatched in oxygenated salt water (36g/L). All fish were maintained in a standard fish water medium (pH 7, 700 µS/cm, containing 150 mg/L CaCO3 and 60 mg/L NaCl) at 28°C with a 12h/12h day-light cycle.

### Breeding procedure

In the late afternoon (4:00 PM, 1h after their last meal) one male and one female were placed in mating tanks (1L) equipped with dividers and filled with standard breeding water (described above), incorporating a few glass marbles for enrichment. At the onset of light (around 8:30 AM), dividers are removed from the tanks. Fish are then reintroduced to the breeding system at the end of the day (6:00 PM), and eggs are collected, rinsed with osmosis water, and transferred to a Petri dish field with standard rearing water.

### F0 reproduction monitoring – Experiment 1

Embryos from the F0 were collected at stage 1 from 4 different wild type strain (AB line(Stuart et al. 1990, Brown et al. 2015)) siblings and 1nl of an oncogenic plasmid (n=376, Hsp70 RAS v12 GFP+ transposase) was injected using a micro injector under a stereomicroscope following (Bambino and Chu 2017). A heat shock was applied at 38°C 24 hours post-injection to activate plasmid expression. Injected embryos were hatched in petri dishes and their fluorescence was observed at 48 hours post-fertilization (hpf) to identify and select embryos expressing the oncogenic gene. Juveniles were then transferred to adult plastic fish tanks at a density varying from 20 to 50 individuals depending on their size. Sex identification was performed at 13 weeks based on visual screening. The emergence of tumors on cancerous fish was observed daily with the method described in (Sophie Tissot, Lena Guimard, Jordan Meliani, Justine Boutry, Antoine M. Dujon, Jean-Pascal Capp, Jácint Tökölyi, Peter A. Biro, Christa Beckmann, Laura Fontenille, Nam Do Khoa, Rodrigo Hamede, Benjamin Roche, Beata Ujvari 2023).

### F1 reproduction monitoring – Experiment 2

Tumoral females were isolated from the F0 generation to mate with wild-type males from the AB strain and generate an F1 generation raised in the same conditions as described above. Then the total number of eggs produced and the sex ratio of each of the F1 fish was observed at week 13^th^. The complete experimental design is represented in the Figure 1A.

**Figure 1:**
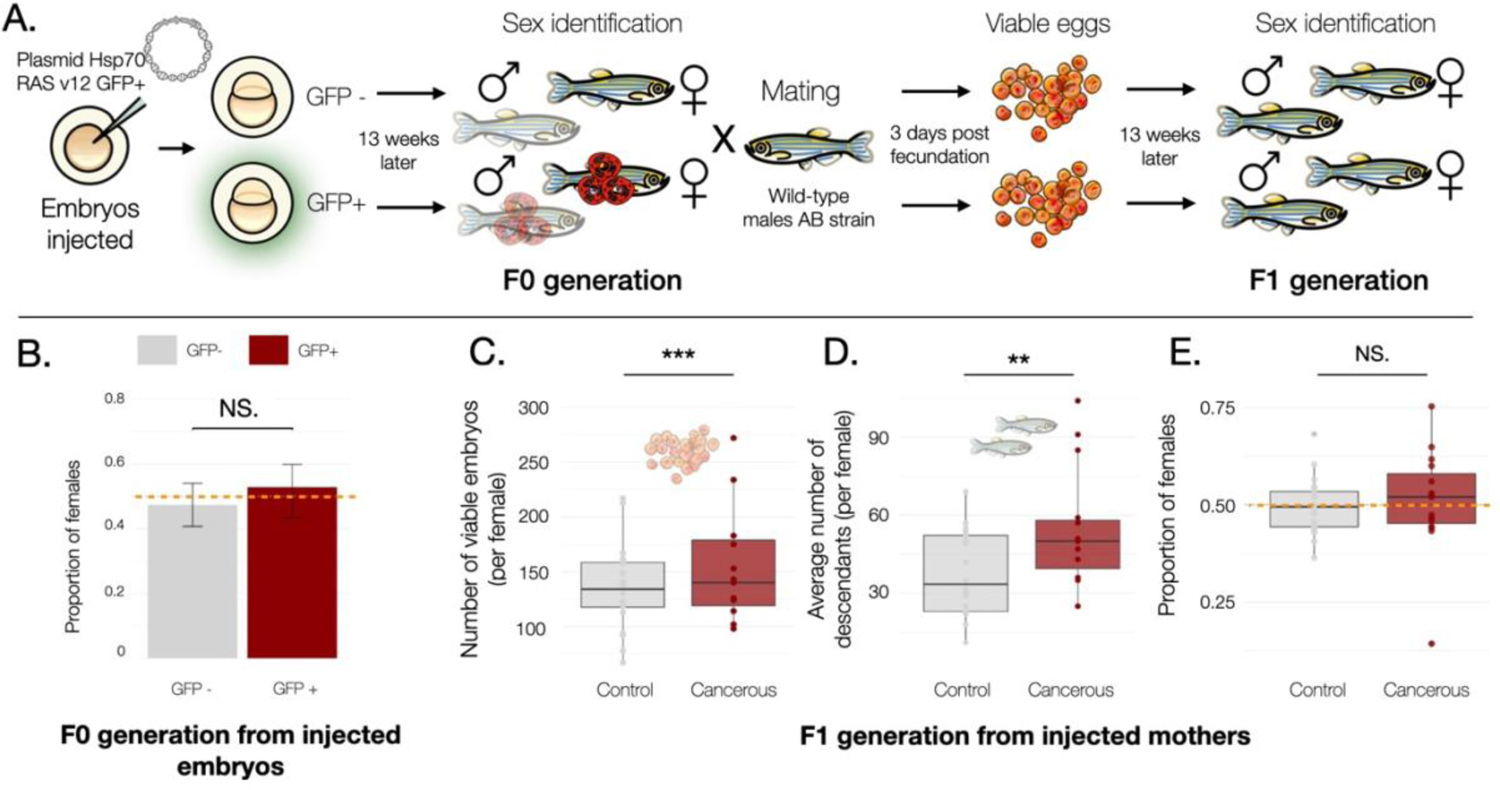
Monitoring the sex ratio of two successive generations after inducing skin cancer in Zebrafish (*Danio rerio*). (A) Experimental design: Embryos were injected with a carcinogenic plasmid. After 48 hours, the inclusion of the plasmid was screened through GFP fluorescence observations. Skin cancer was only observed among the GFP+ group. The individuals emerging from those embryos (F0) were sorted by sex at the 13th week and females were mated with wild-type males. The number of embryos obtained from the female fish was counted 3 days after fecundation (dpf), and the sex of the individuals emerging from those embryos (F1) was identified at the 13th week. (B) The proportion of female fish in the F0 generation injected with an oncogenic plasmid was determined based on whether the plasmid was expressed (GFP+ group in red on the right, n=149) or not (GFP-group in grey on the left, n=228). There was no significant difference in the proportion of females between the two groups (Newcomb proportion test, X-squared = 0.508, df = 1, p-value = 0.476). (C) The number of viable embryos (3 dpf) in the F1 generation of fish was recorded based on whether the mother was cancerous (positive group in red on the right, n=15) or not (negative group in grey on the left, n=20). (D) The average number of descendants per female in the F1 generation of fish at the 13th week, categorized by whether the mother was cancerous (positive group in red on the right, n=15) or not (negative group in grey on the left, n=20). (E) Proportion of female F1 fish generation based on whether the mother was cancerous (GFP+ group in red on the right, n=15) or not (GFP-group in grey on the left, n=20).

### Statistical analyses

#### Statistical Analysis

The sex ratios within the F0 and F1 generations were compared employing binomial distribution, utilizing Generalized Mixed Effect Models (GLMMs). To identify the most pertinent models, we undertook a two-step selection process based on the conditionals Akaike Information Criterion (AICc) and associated weights [detailed in (Zuur et al. 2009)]. First, models were compared incorporating potential combinations of random effects, including the common aquarium, experimental batch for the F0, the breeding date and the experimental batch for the F1. Subsequently, models were evaluated considering the fixed effects of plasmid injection, as well as the number of embryos and of mature adults in the pond at the 13th week for the F0. Analyses were conducted under R© (R version 4.3.1) through the interface Rstudio© (Version 2023.12.0+369). Data files were imported using the readr package (Wickham et al. 2023), while graphical exploration of the data was conducted using GGally (Schloerke et al. 2021). Statistical models were primarily fitted using the glmmTMB package (Brooks et al. 2017) for generalized mixed-effects models. Results were visually presented using the sjPlot package (Lüdecke 2023) to generate model plots. Model selection criteria were applied with the MuMIn package (Bartoń 2023) used for AIC calculation. The equilibrium and dispersion of residuals were evaluated using the DHARMa package (Team et al. 2021), ensuring the robustness and appropriateness of the selected models. Additional analyses were carried out using the MASS package (Venables and Ripley 2002) for supplementary statistical functions. Graphs were generated using ggplot2 (Schloerke et al. 2021) and the software keynotes ©.

## RESULTS

### Impact of Skin Tumors on the Sex Ratio (F0 Generation)

Out of the 376 embryos injected with the oncogenic plasmid surviving until the 13th week, 60.6% (n=228) effectively expressed the plasmid 48 hours later. No significant difference in the sex ratios of zebrafish in the F0 generation was observed between those expressing the oncogenic plasmid (GFP+, Figure 1.B, *p*^=0.52 [0.4338; 0.60], n=228) and the control group (GFP-, Figure 1.B, *p*^==0.48 [0.401; 0.54], n=149) (See supplementary Table S1 and Figure 1.B). Among the 77 female fish incorporating the plasmid and surviving for 13 weeks, 15 developed tumors at the adult stage and were used as mothers for the subsequent experiment.

### Impact of Skin Tumors on the Survival and Sex Ratio of the Offspring (F1 Generation)

After mating with wild-type males, cancerous females (GFP+) produced more viable embryos (Figure 1.C, µ^=159.8 [124.0; 195.6]) compared to control females (GFP-, µ^=135.6 [117.4; 153.8]). The incidence rate ratio (IRR) was 1.26 [1.18 – 1.34], indicating that, on average, a cancerous female produced 1.26 viable embryos for every one produced by a non-cancerous female, with random variations observed across mating dates (See supplementary Table S2 and Figure 1.C, p < 0.001).

Viable embryos from cancerous mothers were more likely to survive until their 13th week (Figure 1.D, µ^=53.9 [41.1; 66.7]) compared to those born from control mothers (GFP-, µ^=37.6 [29.9; 45.2]). The incidence rate ratio (IRR) was 1.19 [1.05 – 1.35], indicating a 1.19 times higher likelihood for embryos from cancerous mothers to reach maturity by the 13th week, with random variations observed across mating dates and experimental batches (See supplementary Table S3 and Figure 1.D, p = 0.005).

Regarding the sex ratio of the F1 generation of fish, statistical model selection revealed no significant influence of the cancerous state of the mother, the number of mature and viable embryos from the same clutch, and no random effect of different mating dates and experimental batches (See supplementary material, Table S3 and Figure 1.E).

## DISCUSSION

The study of the impact of oncogenic processes on the evolutionary ecology of wildlife is a rapidly expanding area of research. However, our understanding is currently limited, as we are far from having explored the full spectrum of biological traits that could potentially be influenced by malignant progression (Thomas et al. 2017).

Chemical pollution in ecosystems has been shown to influence the sex ratio in wild animal populations, particularly in species whose sex is determined by the environment (Decourten and Brander 2017)(Miracle et al. 2011)(Mikó et al. 2020). For instance in the zebrafish, exposure to clotrimazole (an endocrine disrupting chemical), induces male-skewed sex ratios (Brown et al. 2015). Interpretation of the causes responsible for these skews is often complex because multiple factors may act together, including the mode of actions of chemicals, temperature or other environmental parameters such as stress or inbreeding (Dang and Kienzler 2019)(Dang 2016). Thus, the extent to which diseases resulting from pollution, such as cancer, directly impact the sex ratio in populations remains uncertain. Our findings do not support the hypothesis that cancer progression itself is involved in sex ratio variation, neither directly nor through an adaptive response from the host. Indeed, regarding Experiment 1, cancer does not appear to be associated with differential sex-specific mortality at the embryonic stage, nor does it play a role in sex determination (since both phenomena would have skewed the sex-ratio). These results contrasts with numerous studies that have identified sex-specific responses in zebrafish regarding various factors (e.g., (Zhou et al. 2010, Li et al. 2019, Giommi et al. 2022, Lin et al. 2022, Park et al. 2022, van Gelderen et al. 2022, Zhai et al. 2022, Zheng et al. 2022, Zhong et al. 2023)), including cancers (e.g., (Lee et al. 2018, 2021, Mensah et al. 2019, Montal et al. 2023)). One possible explanation is that the cancer type used in our study may have too low aggressiveness, as possibly evidenced by the ability of individuals affected at the embryo stage to develop into adulthood. This study would, therefore, need to be repeated with a more aggressive cancer to potentially illustrate a difference in mortality between the sexes.

An alternative explanation for the absence of cancer influence on sex ratio is that sex ratios are being adjusted in relation to factors such as growth. In many fish, such as Atlantic silversides (*Menidia menidia*), larger females produce more eggs than smaller females while fecundity of males is less dependent on body size because they do not typically monopolize mating opportunities (Conover and Kynard 1981). Thus, the fitness benefit of larger size is greater for females than for males. Growth-dependent sex determination may maximize reproduction in zebrafish in the same way, where faster-growing individuals developing as females and slower-growing individual as males (Lawrence et al. 2008). If tumors impede organismal growth, as observed for intestinal tumor in zebrafish (Enya et al. 2018), this should select for sex ratio adjustment in the opposite direction (towards males) in contrast to that predicted by sex differential embryonic mortality (towards females, [18]). Thus, the predicted direction of sex ratio adjustment may depend on the relative importance of these two opposing forces and could lead to selection for no sex ratio adjustments. Further work with a larger sample size is needed to follow up on this study, examining the consequences of skin tumors on growth rates and the condition factor of fish. Additionally, it would be interesting to investigate the influence of inter-individual differences in skin tumor development on our observations.

Considering the second experiment, our results show that there is no significant difference in the sex ratio of broods from cancerous and non-cancerous females. Although many fish species now develop cancers in the wild (Baines et al. 2021b), no study has explored the links between this pathology and egg or sperm quality. Further studies would be necessary to clarify this point. An intriguing pattern observed in this study, consistent with prior research on the reproductive biology of organisms bearing tumors ((Arnal et al. 2017)(Boutry et al. 2022b)(Jones et al. 2008)), is that females with cancer appear to yield more offspring and with a survival until sexual maturity that is enhanced. This supports the idea of a greater allocation of reproductive resources in females whose life expectancy is compromised by cancer progression. However, further studies on the quality of eggs produced by both cancerous and non-cancerous females would be necessary to provide additional information on maternal investment in eggs. Further data, particularly on other fish/cancer models, would also be needed to establish whether or not generalizations can be made.

In summary, this pioneering study explores the complex responses of an aquatic vertebrate organism to oncogenic processes at different life stages. The influence of skin cancer on the reproductive ecology of zebrafish does not support the idea that disease progression interferes with the mechanisms governing sex-ratio variations in this species, either directly or through adaptive responses. Nevertheless, the results are consistent with previous studies suggesting that organisms that develop cancer tend to maximize their immediate reproductive effort before dying prematurely. This study highlights the eco-evolutionary dynamics of the responses developed by organisms increasingly confronted with oncogenic compounds in contemporary ecosystems polluted by anthropogenic activities.

## Supporting information

Data of the F0 fish

Data of the F1 fish

## Acknowledgments

This work was funded by the CNRS (IRP CANECEV), the Hoffmann Family, and by the following grant: EVOSEXCAN project (ANR-23-CE13-0007).

## SUPPLEMENTARY TABLES

**Table S1:** Comparisons of the results of different generalized models explaining the sex-ratio in the F0 generation (binomial law) of Zebrafish (*Danio rerio*). Models with variations in their random or fixed effect are compared using degrees of freedom, AICc values and weights.

| Random model selection | Df | AICc | Weights |
| --- | --- | --- | --- |
| Sex ratio ~ GFP * Total number of individual + (1\|Exp/Aquarium) | 6 | 87.38 | 0.009 |
| Sex ratio ~ GFP * Total number of individual + (1\|Aquarium) | 5 | 82.43 | 0.112 |
| Sex ratio ~ GFP * Total number of individual | 4 | 78.32 | 0.879 |
| Fixed model selection | Df | AICc | Weights |
| Sex ratio ~ GFP * Total number of individual | 2 | 91.39 | 0.000 |
| Sex ratio ~ GFP + Total number of individual | 4 | 85.56 | 0.003 |
| Sex ratio ~ GFP | 3 | 77.84 | 0.121 |
| Sex ratio ~ Total number of individual | 3 | 81.90 | 0.016 |
| Sex ratio ~ 1 | 5 | 73.91 | 0.864 |

**Table S2:** Comparisons of the different generalized model explaining the survival rate in the F1 generation (binomial law) of Zebrafish (*Danio rerio*). Models with variations in their random or fixed effect are compared using degrees of freedom, AICc values and weights. The estimator of the fixed and random effect of the best model is shown in the last part of the table, with their associated incidence rate ratio (IRR), confidence interval (CI), p-value, number of random categories, residual variance (σ^2^), marginal and conditional correlation coefficient (R^2^).

| Random model selection | Df | AICc | Weights |
| --- | --- | --- | --- |
| Viable embryo ~ GFP + (1\|Exp/Mating date) | 4 | 647.67 | 0.218 |
| Viable embryo ~ GFP + (1\|Mating date) | 3 | 645.12 | 0.782 |
| Viable embryo ~ GFP | 2 | 786.12 | 0.000 |
| Fixed model selection | Df | AICc | Weights |
| Viable embryo ~ GFP + (1\|Mating date) | 2 | 653.53 | 1.000 |
| Viable embryo ~ 1 + (1\|Mating date) | 3 | 698.04 | 0.000 |
| Final best model | IRR | CI | P-Value |
| GFP [pos] | 1.26 | 1.18 – 1.34 | <0.001 |
| Random Effects |  |  |  |
| $\sigma^2$ | 0.01 | | |
| Nb mating date | 11 |  |  |
| Marginal R2 / Conditional R2 | 0.186 / 0.905 |  |  |

**Table S3:** Comparisons of the different generalized model explaining the sex-ratio in the F1 generation (binomial law) of Zebrafish (*Danio rerio*). Models with variations in their random or fixed effect are compared using degrees of freedom, AICc values and weights. The estimator of the fixed and random effect of the best model is shown in the last part of the table, with their associated incidence rate ratio (IRR), confidence interval (CI), p-value, number of categories, residual variance (σ^2^), marginal and conditional correlation coefficient (R^2^).

| Random model selection | Df | AICc | Weights |
| --- | --- | --- | --- |
| Mature fish / Viable embryo ~ GFP + (1\|Exp/Mating date) | 4 | 332.14 | 0.949 |
| Mature fish / Viable embryo ~ GFP + (1\|Mating date) | 3 | 329.58 | 0.051 |
| Mature fish /Viable embryo ~ GFP | 2 | 335.71 | 0.000 |
| Fixed model selection | Df | AICc | Weights |
| Mature fish / Viable embryo ~ GFP + (1\|Exp/Mating date) | 4 | 339.96 | 0.668 |
| Mature fish / Viable embryo ~ 1 + (1\|Exp/Mating date) | 3 | 341.36 | 0.332 |
| Final best model | IRR | CI | P-Value |
| GFP [pos] | 1.19 | 1.05 – 1.35 | 0.005 |
| Random Effects |  |  |  |
| $\sigma^2$ | 3.29 | | |
| Nb mating date | 11 |  |  |
| Nb experimental batch | 2 |  |  |
| Marginal R2 / Conditional R2 | 0.186 / 0.905 |  |  |

**Table S4:** Comparisons of the results of different generalized models explaining the sex-ratio in the F1 generation (binomial law) of Zebrafish (*Danio rerio*), their degrees of freedom, AICc values and weights.

| Random model selection | Df | AICc | Weights |
| --- | --- | --- | --- |
| Sex ratio ~ GFP + Nb Mature fish + Viable embryo + (1\|Exp/Matng date) | 6 | 193.06 | 0.045 |
| Sex ratio ~ GFP + Nb Mature fish + Viable embryo + (1\|Mating date) | 5 | 190.12 | 0.194 |
| Sex ratio ~ GFP + Nb Mature fish + Viable embryo GFP | 4 | 187.39 | 0.761 |
| Fixed model selection | Df | AICc | Weights |
| Sex ratio ~ GFP x Nb Mature fish x Viable embryo GFP | 8 | 281.23 | 0.000 |
| Sex ratio ~ GFP + Nb Mature fish x Viable embryo GFP | 5 | 238.13 | 0.000 |
| Sex ratio ~ GFP x Nb Mature fish + Viable embryo GFP | 5 | 227.75 | 0.000 |
| Sex ratio ~ GFP x Viable embryo GFP + Nb Mature fish | 5 | 229.54 | 0.000 |
| Sex ratio ~ Nb Mature fish x Viable embryo GFP | 4 | 232.58 | 0.000 |
| Sex ratio ~ GFP x Nb Mature fish | 4 | 212.94 | 0.000 |
| Sex ratio ~ GFP x Viable embryo GFP | 4 | 217.95 | 0.000 |
| Sex ratio ~ Nb Mature fish + Viable embryo GFP | 3 | 211.40 | 0.000 |
| Sex ratio ~ GFP + Nb Mature fish | 3 | 202.05 | 0.000 |
| Sex ratio ~ GFP + Viable embryo GFP | 3 | 204.98 | 0.000 |
| Sex ratio ~ GFP + Nb Mature fish + Viable embryo GFP | 4 | 216.77 | 0.000 |
| Sex ratio ~ Nb viable embryo GFP | 2 | 199.89 | 0.001 |
| Sex ratio ~ GFP | 2 | 190.49 | 0.082 |
| Sex ratio ~ Nb Mature fish | 2 | 196.84 | 0.003 |
| Sex ratio ~ 1 | 1 | 185.66 | 0.913 |

## Notes

### Competing Interest Statement

The authors have declared no competing interest.

